# Metapopulation Structure of Diatom-associated Marine Bacteria

**DOI:** 10.1101/2021.03.10.434754

**Authors:** Liping Qu, Xiaoyuan Feng, Yuerong Chen, Lingyu Li, Xiaojun Wang, Zhong Hu, Hui Wang, Haiwei Luo

## Abstract

Marine bacteria-phytoplankton interaction ultimately shapes ecosystem productivity. The biochemical mechanisms underlying their interactions become increasingly known, yet how these ubiquitous interactions drive bacterial evolution has not been illustrated. Here, we sequenced genomes of 294 bacterial isolates associated with 19 coexisting diatom cells. These bacteria constitute eight genetically monomorphic populations of the globally abundant Roseobacter group. Six of these populations are members of *Sulfitobacter*, arguably the most prevalent bacteria associated with marine diatoms. A key finding is that populations varying at the intra-specific level have been differentiated and each are either associated with a single diatom host or with multiple hosts not overlapping with those of other populations. These closely related populations further show functional differentiation; they differ in motility phenotype and they harbor distinct types of secretion systems with implication for mediating organismal interactions. This interesting host-dependent population structure is even evident for demes within a genetically monomorphic population but each associated with a distinct diatom cell, as shown by a greater similarity in genome content between isolates from the same host compared to those from different hosts. Importantly, the intra- and inter-population differentiation pattern remains when the analyses are restricted to isolates from intra-specific diatom hosts, ruling out distinct selective pressures and instead suggesting coexisting microalgal cells as physical barriers of bacterial gene flow. Taken together, microalgae-associated bacteria display a unique microscale metapopulation structure, which consists of numerous small populations whose evolution is driven by random genetic drift.

---

Since marine phytoplankton contribute one-half of global primary production (*1*) and since heterotrophic bacterioplankton process 40-50% of the carbon fixed by marine phytoplankton (*2, 3*), bacteria-phytoplankton interaction is an important process that ultimately drives carbon cycling and regulates ecosystem productivity. The physical interface mediating these ubiquitous interactions is a microzone of a few cell diameters immediately surrounding an individual phytoplankton cell, which is termed as ‘phycosphere’ (*4*). In the phycosphere of eukaryotic marine phytoplankton lineages (e.g., diatoms, dinoflagellates, coccolithophores, and green pico-algae), bacterial communities are consistently dominated by a handful of taxa including *Rhodobacteraceae* (mostly the Roseobacter group), *Alteromonadaceae*, and *Flavobacteriaceae* (*4, 5*), and the bacterial community assembly at the phycosphere of a given phytoplankton species is reproducible (*6*). These recurrent patterns in part result from the innate ability of phytoplankton to modulate their bacteria consortia by secreting secondary metabolites such as rosmarinic acid and azelaic acid released by a diatom species, which promote the attachment and growth of certain roseobacters but suppress opportunistic bacteria (*7*). Another important mechanism is the resource-based niche partitioning among these major bacterial associates at nutrient-enhanced phycosphere (*8, 9*). For example, diatoms may use their abundant metabolites such as 2,3-dihydroxypropane-1-sulfonate (DHPS) for targeted feeding of beneficial symbionts among which roseobacters represent a dominant group (*10*).

Recent studies have revealed a greater diversity of the mechanisms underlying the symbiosis between Roseobacter lineages and phytoplankton species than previously appreciated. Some roseobacters such as *Sulfitobacter* spp., *Ruegeria* spp. and *Dinoroseobacter* spp. establish mutualistic interactions with diatoms and microscopic green algae by providing growth factors such as vitamins and indole-3-acetic acid (IAA) to phytoplankton hosts in exchange for labile organic matter (*10–12*), whereas others such as *Sulfitobacter* spp. are virulent to coccolithophores by releasing algicides (*13*). In another type of interaction, some roseobacters including *Phaeobacter* spp. and *Dinoroseobacter* spp. each act initially as a mutualist and later as a parasite of coccolithophores and dinoflagellates, respectively (*14, 15*). An important implication from these studies is that closely related roseobacters (e.g., members of *Sulfitobacter*) may employ different mechanisms to interact with phytoplankton. This high diversity of roseobacter-phytoplankton interaction therefore suggests that the phycosphere may act as an effective barrier of gene flow among symbiotic roseobacters associated with different phytoplankton cells, leading to independent evolution of even closely related roseobacter populations in the seemingly well mixed seawater.

To test this hypothesis, we sought to determine the population structure of roseobacters colonizing the phycosphere of coexisting microalgal cells. A few environmental factors including nutrient availability (*16*), interactions with phages (*17*) and phytoplankton or particles (*18*) are known to drive roseobacter population differentiation. To single out diatom phycosphere from other confounding factors that may drive roseobacter evolution, populations associated with diatoms isolated from a single seawater sample were analyzed. We collected 1 L of seawater from the Pearl River Estuary located at the northern boundary of the South China Sea, isolated 45 diatom cells varying at the level of phylogenetic relatedness (Fig. 1A, Table S1), cultivated over 850 roseobacters associated with 19 of these diatoms (Fig. 1A, Table S2), and sequenced 294 genomes of these roseobacters (Table S3), among which six genomes are complete and closed (Table S4) by additional sequencing with Nanopore (Supplemental Text 1.1–1.3). These newly sequenced roseobacters comprise eight clades (Fig. 1B; see methods in Supplemental Text 1.4), among which six are related to three species of *Sulfitobacter*, a roseobacter genus most commonly found on diatoms (*19*). These include clade-2a and clade-2b related to *Sulfitobacter pseudonitzschiae*, clade-2c related to *S. geojensis*, clade-2d, clade-2e1 and clade-2e2 related to *S. mediterraneus* (Fig. 1B). The remaining clade-1 and clade-3 are related to *Marivita cryptomonadis* and *Ponticoccus* sp. LZ-14, respectively, which are distantly related to *Sulfitobacter*. Members within each clade share identical 16S rRNA genes, display whole-genome average nucleotide identity (ANI) over 99.99% (Fig. S1), and vary up to 45 non-singleton single nucleotide polymorphisms (SNPs) in which the rare variant occurs in at least two genomes (Table S5). Despite this genetic monomorphism, there is ample evidence that the within-clade members are predominantly from the environment rather than a result of clonal replications during the laboratory cultivation of the diatom cells, a required process prior to roseobacter isolation (see Supplemental Text 2, Fig. S2 and Table S5).

**Fig 1.**
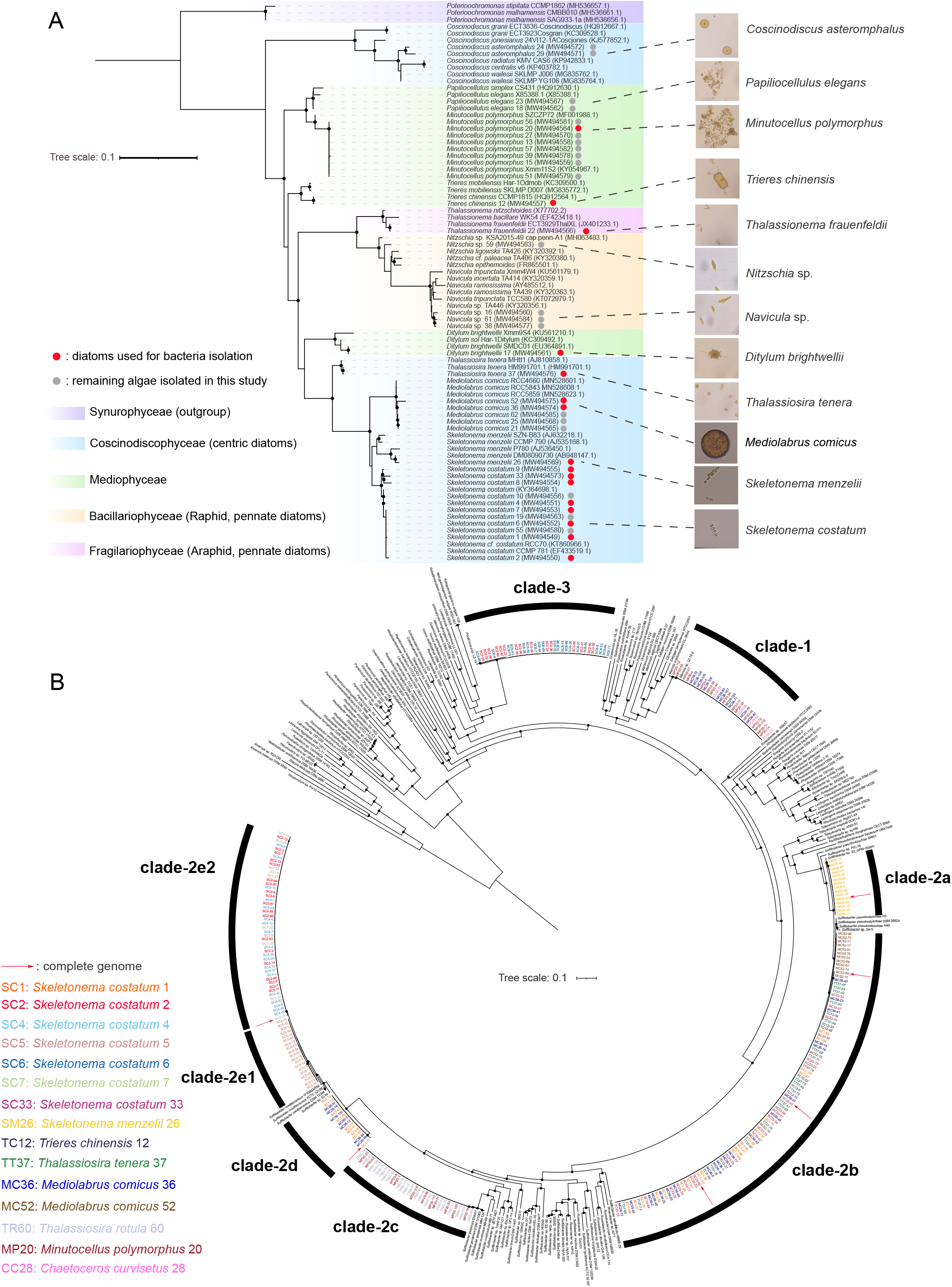
Phylogenetics of diatoms and Roseobacters isolated from this study. (A) Maximum likelihood (ML) tree showing the phylogenetic positions of diatoms isolated in this study. The phylogeny was inferred using IQ-TREE based on the 18S rRNA gene sequences with lengths longer than 1,600 bp. Three *Synurophyceae* strains were used as the outgroup. Solid circles in the phylogeny indicate nodes with ultrafast bootstrap (UFBoot) values > 95%. Diatom strains used for bacteria cultivation and other isolated microalgae are marked with red and gray dots, respectively. The photos of microalgae obtained under an optical microscope are shown on the right panel. (B) ML phylogenomic tree showing the phylogenetic positions of roseobacters isolated from this study. The phylogeny was inferred using IQ-TREE based on the concatenation of 120 conserved bacterial proteins (see methods). Solid circles in the phylogeny indicate nodes with UFBoot values > 95%. The associated microalgae of bacterial strains are differentiated with colors. The six complete and closed genomes are marked with red arrows.

A key finding is that closely related roseobacter populations associated with different diatom cells are often genetically differentiated (see methods in Supplemental Text 1.6). This is clearly supported by populations diverged at different phylogenetic depths. The two clades (clade-2a and clade-2b) related to *S. pseudonitzschiae* share 94.16% ANI (Fig. S1), which is at the boundary (95% ANI) delineating a distinct species (*20*). The SNP density within each clade is extremely low across the whole genome but becomes very high between the clades (Fig. S3), indicating that these two clades each have fixed distinct alleles and are thus genetically isolated. The clade-2a is composed solely of members isolated from *Skeletonema menzelii* 26 (abbreviated as ‘SM26’), whereas the clade-2b comprise members from five diatom cells of the class *Coscinodiscophyceae* affiliated with three species including *Skeletonema costatum* (SC1 & SC33), *Mediolabrus comicus* (MC36 & MC52), and *Thalassiosira tenera* (TT37), and one diatom cell of a distantly related class *Mediophyceae* affiliated with the species *Trieres chinensis* (TC12) (Fig. 1B, Table S2). Among the three clades related to *S. mediterraneus*, clade-2e1 and clade-2e2 share 97.78% ANI, suggesting that they have not yet separated into two distinct species. However, a dramatic increase in SNP density of between-clade comparisons compared to the within-clade comparisons across the whole genome (Fig. S4A,B,C) indicates that these two clades are under ongoing speciation. The clade-2d is more divergent, showing 89.34% ANI to clade-2e1 and 89.51% ANI to clade-2e2, though it shares with the latter two clades very high similarities (99.86% and 99.93%, respectively) at the 16S rRNA gene sequence. The SNP density derived from the comparison between clade-2e1 and clade-2e2 is much lower than when each is compared to clade-2d (Fig. S4C,D,E), suggesting that the genomes are differentiated to a greater extent when the phylogenetic depth becomes larger for these overall very closely related clades. Importantly, these clades each have a distinct host range. Members of clade-2e1 are exclusively associated with a single diatom host (SC5), whereas members of clade-2e2 are from three hosts of the same diatom species (SC2, SC4 and SC7) and members of clade-2d are from hosts of two different families (SC1 and MC36) (Fig. 1B, Table S2). Population differentiation of the sampled clades is further supported by genome rearrangement in both chromosome and plasmids, as shown by the more conserved gene order of within-clade members (Fig. S2) compared to that of between-clade members (Fig. S5) for clade-2a and clade-2b. Multiple genome rearrangement events were also observed between clade-2d and clade-2e2 (Fig. S6), though within-clade comparisons cannot be made because no more closed genomes are available in both clades.

Diatom-dependent differentiation is also evident from the more closely related roseobacter demes associated with different hosts but sharing membership of the same clade. Because of the genetic monomorphism at the core genomes (Table S5), the within-clade members do not show a reliable phylogenetic structure. We therefore turned to explore the accessory genes which are shared by a subset of the genomes under comparison. A simple clustering based on the presence and absence pattern of the accessory genes identified clusters corresponding to distinct diatom hosts. In clade-2b, for example, members associated with SC33 largely constitute an independent cluster separated from members associated with other hosts (Fig. 2A). Likewise, in clade-2c, members from *Minutocellus polymorphus* 20 (MP20) are overall well separated from those associated with *Thalassiosira rotula* 60 (TR60) (Fig. 2B); in clade-2e2, most members associated with SC2 are separated from those with other hosts (Fig. 2C); and in clade-3, members from SC2 and those from SC6 are generally clustered into two separate groups (Fig. 2D). For the remaining clades (clade-2d & clade-1) with members associated with multiple hosts, host-dependent clustering is not obvious (Fig. S7).

**Fig. 2.**
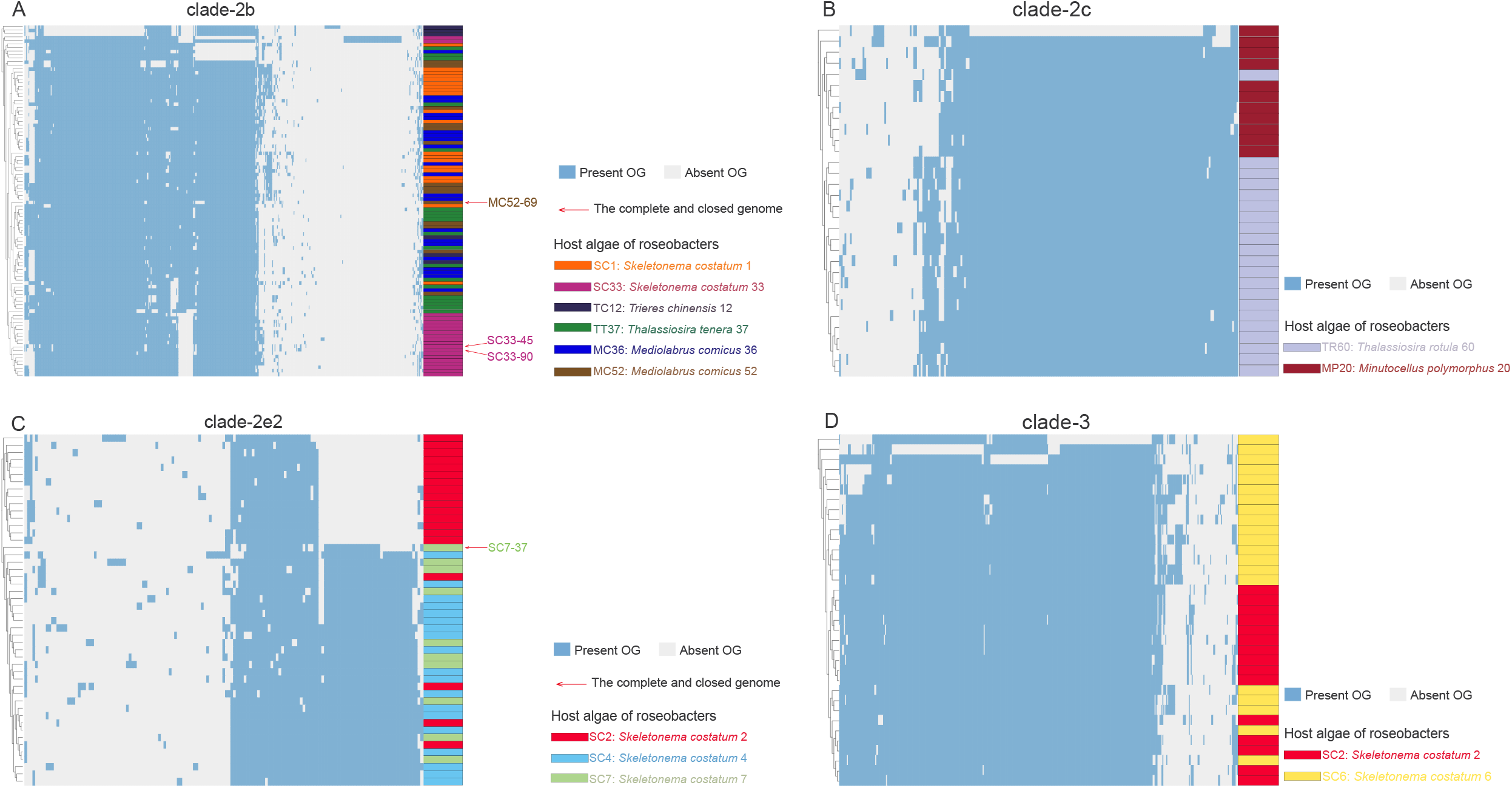
The clustering of accessory genes in the genomes of (A) clade-2b, (B) clade-2c, (C) clade-2e2 and (D) clade-3. The dendrogram of genome clustering was generated based on the presence and absence of their orthologous gene families (OGs), which are colored in blue and gray, respectively. The associated microalgae of bacterial strains are differentiated with colors. The complete and closed genomes are marked with red arrows.

We further identified important evidence for population differentiation at the functional level. The secretion systems are well known to mediate bacteria-bacteria and bacteria-host interactions. Interestingly, the presence and absence pattern of three secretion systems differentiates the clade-2e1, clade-2e2 and clade-2d related to *S. mediterraneus* (Table S6). Specifically, the type VI secretion system (T6SS) transports effector proteins into both prokaryotic and eukaryotic cells in a contact-dependent manner (*21, 22*). This system was reported in only a few roseobacter lineages (*23, 24*). We showed an exclusive presence of a T6SS gene cluster on the chromosome of clade-2d members. Another uncommon secretion system in roseobacters, thus far only reported in the roseobacter species *Marinovum algicola* (*25*), is the type II secretion system (T2SS), which promotes the release of folded proteins, mainly extracellular enzymes such as proteases, lipases, phosphatases, and polysaccharide hydrolases, to the extracellular milieu or displayed on the cell surface (*26*). We found an exclusive occurrence of a T2SS cluster on a plasmid of the clade-2d members. In terms of the type IV secretion system (T4SS), the *virB/D4* type secretes effector proteins and plasmid DNA to target both bacteria and hosts (*27–29*), whereas the *trb* type transports plasmid DNA between bacteria (*29*). While the *virB/D4* is commonly found among roseobacters (*30*), the *trb* is rare in these bacteria. Consistent with this pattern, a *virB/D4* gene cluster was found in all three clades, but a *trb* gene cluster was exclusively identified on the chromosome of clade-2e2 members. In the case of the *S. pseudonitzschiae* related clades, both clade-2a and clade-2b carry the *virB/D4*-based T4SS, but they differ in copy numbers. The *virB/D4* copy number difference was similarly found between the three *S. mediterraneus* related clades, but a unique observation was that the clade-2d members possess an additional copy on their chromosomes instead of the plasmids where this type of secretion system usually locates. No other secretion systems were found in clade-2a and clade-2b. Gene clusters encoding all secretion systems locate within the genomic islands except the T6SS of clade-2d (Table S6), suggesting that roseobacter-diatom and/or roseobacter-bacteria interactions are highly dynamic.

Since these secretion systems may mediate either pathogenic, or commensal, or mutualistic relationships with hosts and/or other bacteria (*21, 26, 28, 29, 31, 32*), their differential presence among the clades suggests that distinct clades may exert different and even opposite physiological effects on the diatom hosts. This motivated us to set up experimental assays (Supplemental Text 1.7) to compare the effects of co-culture of diatom and roseobacter, the latter represented by each of the *S. mediterraneus* and *S. pseudonitzschiae* clades, on the growth of the diatom. Among the 11 tested roseobacter isolates, eight significantly promoted the growth of the diatom, whereas the remaining three did not significantly change the growth rate of the diatom (Fig. S8). We did not observe consistent differences between closely related clades regarding their effects on the diatom growth (Fig. S8). While this assay was motivated by the observation of clade-specific secretion systems, there is no direct link between the algal growth change and the differential presence of the secretion systems in the bacterial symbionts.

Another important metabolic trait relevant to roseobacter-phytoplankton interaction is the bacterial motility (*33*), which is differentially present among these related roseobacter clades. Three phylogenetically distinct flagellar gene clusters (FGCs) designated as *fla1*, *fla2* and *fla3* (Fig. 3A) have been identified in the Roseobacter group, and carrying any of them may enable motility (*34, 35*). Among these, *fla2* is present in a plasmid of the *S. pseudonitzschiae* related clade-2e1, clade-2e2 and clade 2d, whereas *fla1* was exclusively found on the chromosome of clade-2e2 (Fig. 3A, Table S6; see methods in Supplemental Text 1.8). Despite the presence of the flagellum-encoding gene clusters, the flagella were not detected by transmission electron microscopy (Fig. 3B) and the motility phenotype was not observed under the experimental condition (Fig. 3B; Supplemental Text 1.9). The lack of flagella and motility in these related clades is likely due to inappropriate physicochemical conditions set in the laboratory experiment, as temperature (*36*), pH (*37*), salinity (*38*) and metal ions (*39*) were demonstrated to induce the expression of the FGC genes in other bacteria. In terms of the *S. pseudonitzschiae* related clade-2a and clade-2b, they did not possess any type of FGC (Table S6), but both instead carry homologs of two candidate gene clusters (i.e., type-IVb tight adherence pilus gene cluster; Table S6) recently hypothesized to be responsible for dendritic motility (*34*). We showed that the clade-2b members possess an additional copy located within a genomic island compared to clade-2a members, consistent with the greater swimming and dendritic motilities observed in the former (Fig. 3C).

**Fig. 3.**
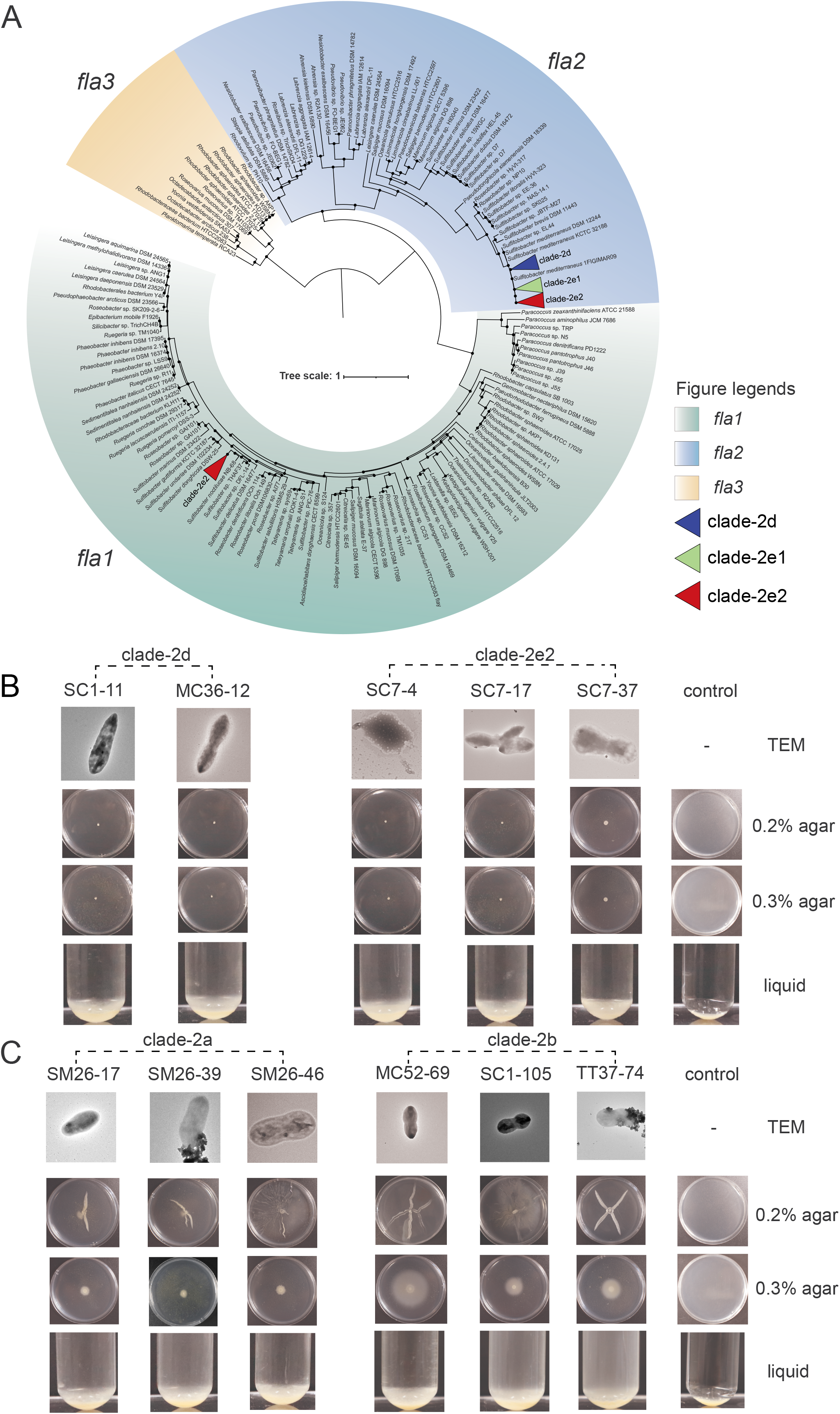
The metabolic traits. (A) ML phylogenetic tree of the three homologous types of flagellar gene clusters (FGCs) found in the Roseobacter group. This phylogeny was built based on four marker flagellar proteins (FliF, FlgI, FlgH and FlhA). Solid circles in the phylogeny indicate nodes with UFBoot values > 95%. (B, C) Photos showing the cellular morphology under a transmission electron microscope (TEM), motility on the 2216E agar plates with 0.2% or 0.3% agar, and sedimentation phenotypes in the liquid 2216E medium. Eleven representative strains were used in the assay, including two from clade-2d (B), three from clade-2e2 (B), three from clade-2a (C) and three from clade-2b (C). All strains from clade-2e1 were lost.

Previous studies demonstrated that selection for niche adaptation drives sympatric population differentiation in free-living prokaryotic lineages (*18, 40–42*) In the present study, we provided evidence that closely related but genetically discrete populations of several sympatric Roseobacter lineages each have a distinct diatom host range. The pattern of *Sulfitobacter mediterraneus* related populations (clade-2e1, clade-2e2, clade-2d) is of particular interest. This is because populations of this species varying at different stages of differentiation, including those at the very beginning (i.e., within-clade demes differentiated only by accessory gene content), at the middle (i.e., closely related clade under ongoing speciation), and at the completion of speciation, were captured and each found to be associated with different hosts of the same diatom species *Skeletonema costatum*. Since members of the same microalgal species likely release a similar set of organic compounds to the phycosphere and impose other physicochemical parameters (e.g., reactive oxygen species) at similar levels, the observed differentiation of the symbiotic bacterial populations is less likely driven by ecological selection imposed by differential exposure to different microalgal exudates. Instead, symbiotic bacteria may be trapped in the phycosphere (*43*), leading to a reduced opportunity of recombination between bacteria associated with different diatom cells compared to those within the same phycosphere. This is a new mechanism of bacterial population differentiation in the pelagic ocean and represents one of the few examples of population differentiation at the sympatric scale due to physical barriers of gene flow. To put it in context, previous cases of bacterial population differentiation in a sympatric pelagic environment were linked to ecological barriers of gene flow, such as differentiated populations colonizing organic particles of different sizes (*41*) or those inhabiting the bulk seawater versus phycosphere/particles (*18*).

Our observation of subdivision of highly closely related populations each showing genetic monomorphism has important implications for understanding the population structure of the diatom-associated symbiotic roseobacters. Similar population structure was previously demonstrated in obligately host-dependent bacteria such as endosymbionts subjected to repeated bottlenecks during transmissions to new hosts in small numbers of bacterial cells (*44*), and also proposed for generalist marine bacteria such as *Vibrio* spp. which experience short bursts as a result of intensive use of ephemeral resources like organic particles followed by dispersal and colonization of new particles with low numbers of cells (*45*). These two known mechanisms lead to the formation of “metapopulation structure”, in which the population is divided into subpopulations each colonizing a transient resource such as hosts and particles (*45*). Our results suggest that the population structure of the diatom-associated roseobacters aligns well with the metapopulation structure. Hence, phycosphere colonization represents a new mechanism leading to bacterial metapopulation structure in the pelagic ocean. The exact processes leading to the formation of metapopulation structure of these diatom-associated roseobacters remains unknown, however. It could be a result of short burst owing to intensive use of the organic substrates enriched in the phycosphere. It is also possible that bacteria-diatom associations have been maintained at the evolutionary timescale, such that diatom host-dependent population differentiation is evident even at the completion of speciation.

Formation of metapopulation structure in a bacterial species leads to a reduced effective population size (*N_e_*) of the species (*45*), a key parameter in understanding the population genetic mechanism underpinning biological evolution and defined as the size of an ideal population carrying the same amount of the neutral genetic diversity as is observed in the real population (*45, 46*). Because *N_e_* is the inverse of the power of random genetic drift (*45*), the reduced *N_e_* of a bacterial species owing to the formation of metapopulation structure suggests an increased power of genetic drift in driving the evolution of diatom-association roseobacters. As a consequence, the diatom-associated roseobacter populations are predicted to more readily accept the horizontally transferred genetic elements, which are often mildly deleterious owing to the selfish propagation of most mobile genetic elements at the expense of cellular fitness (*47*) but may also carry functional traits such as antimicrobial genes that increase competitive advantages of the bacteria at the phycosphere. Given that diatoms and roseobacters are among the most abundant phytoplankton and bacterial groups, respectively, in today’s ocean (*19*), our findings have important implications for biogeochemical cycles mediated by bacteria-phytoplankton interactions.

It is important to clarify that while the concept of phycosphere has been adopted in this and many other studies, there has been no direct experimental evidence for its occurrence because of technological challenges in separating this microenvironment from the bulk seawater (*4*). Prior studies established microalgal phycosphere as a hotspot of carbon and nutrient cycling in the pelagic ocean. Here, we revealed that phycosphere of diatom cells may act as an effective physical barrier of gene flow between nearly identical symbiotic roseobacters, thereby conferring a new role of phycosphere in driving the evolution of pelagic marine bacteria.

## Acknowledgements

We thank Xingqin Lin for helpful discussion. We also thank Qingyu Li for assisting microalgal isolation and identification, Min Yang for helping with microalgal cell counting, Fumin Yang, Qing Zhang and Huiling Chen for their help in bacteria isolation, Xiaoyu Yang for assisting a preliminary experiment of the motility assays. H.W. was supported by the National Natural Science Foundation of China (41676116), Key Special Project for Introduced Talents Team of Southern Marine Science and Engineering Guangdong Laboratory (Guangzhou) (GML2019ZD0606), and Guangdong Science and Technology Department (2019A1515011139). H.L. was supported by the Hong Kong Research Grants Council General Research Fund (14163917) and the Hong Kong Research Grants Council Area of Excellence Scheme (AoE/M-403/16).

## Author Contributions

H.L. conceptualized the work, designed this study, directed the bioinformatics analyses, interpreted the data, and wrote the main manuscript. H.W. directed the sampling work, diatom isolation, and roseobacter cultivation, experimental assay, and related writing, co-interpreted the data, and provided comments to the manuscript. L.Q. collected the sample, performed cultivation and characterized the cultures. X.F. and L.Q. performed all the bioinformatics, co-interpreted the data, drafted the technical details, and prepared figures and tables. L.L. contributed to bacterial isolation, and L.L. and Y.C. performed physiological assays. X.W. contributed to the bioinformatics. H.Z. contributed to the discussion and provided comments to the manuscript.

## Data availability

The 18S rRNA gene sequences of the diatoms are available at NCBI under the accession number MW494549 - MW494585. The raw genomic sequencing data and assembled genomes of the 294 isolates are available at NCBI under the accession number PRJNA691705.

## Supplementary Materials

### Text 1. Methods

#### 1.1 Isolation, identification and phylogenetic analysis of the microalgae

One-liter of seawater (0.5 m below surface) was collected from the Pearl River Estuary (113.7221° E, 21.9935° N) in January 2018 (*48*), stored in a cooler (4 °C) and transferred to the laboratory within 24 hours. Single cells of microalgae were isolated from the seawater using micropipettes under optical microscope following a previously described method (*49*). The microalgal cells were then washed repeatedly using fresh axenic F/2 medium (*50*) to remove the free-living bacteria and loosely associated bacteria around microalgal cells (*51*). The washed microalgal cells each were inoculated in a 24-well plate containing 1 mL of fresh F/2 medium to increase the cell density. After incubation at 20°C with 12 h/12 h light-dark cycle at 200 μmol photons m^-2^ s^-1^ for 3-5 days, the microalgal culture in each well was transferred into a conical flask with 30 mL of fresh F/2 medium to increase the biomass of microalgae. In the exponential growth phase, 15 mL of the cultures each were fixed with 1% Lugol’s solution and sent to the Fujian Provincial Department of Ocean and Fisheries for morphological characterization and species identification. Besides, the taxonomy of microalgae was further validated using 18S rRNA genes. The DNA was extracted from another 10 mL of the microalgal cultures using the CTAB method (*52*). The 18S rRNA genes were amplified using primers (SSU-F: 5’-ACCTGGTTGATCCTGCCAGT-3’ and SSU-R: 5’-TCACCTACGGAAACCTTGT-3’) following a previous study (*53*) and sent for sequencing using the Sanger platform. These 18S rRNA gene sequences were subjected to a maximum likelihood (ML) phylogenetic analysis along with a few reference sequences, which were selected based on a preliminary BLASTn (*54*) result against the NCBI nt database. These sequences were aligned using Clustal Omega v1.2.4 (*55*) with default parameters and trimmed using trimAl v1.4.rev15 (*56*) with ‘-automated1’ option. The phylogenetic tree was constructed using IQTREE v1.6.12 (*57*) with the Modelfinder (*58*) for model selection, and 1,000 ultrafast bootstrap replicates were sampled to assess the robustness of the phylogeny (*59*). The phylogenetic tree was visualized using iTOL v5.6 (*60*).

#### 1.2 Isolation and identification of bacteria in phycosphere

The 1 mL of microalgal cultures each were collected during the logarithmic growth phase followed by a 10-fold serial dilution. Solid 2216E agar plates (BD Bioscience, USA) were spread with 100 μL of each dilution and incubated at 25 °C for one week. Single colonies were selected and repeatedly streaked on 2216E agar plates to isolate and purify bacterial strains initially associated with microalgae. The purified bacterial strains were subjected to the colony polymerase chain reaction (colony PCR) to retrieve the 16S rRNA genes for taxonomy identification. The 16S rRNA genes were amplified using the universal primers 27F and 1492R as described previously (*61*) and were partially (~700 bp) sequenced using 27F on the Sanger platform at Invitrogen Trading (Shanghai) Co., Ltd. Their taxonomy was inferred using BLASTn on the NCBI website, and a total of 294 strains with top BLAST hit to the Roseobacter group were kept for genome sequencing.

#### 1.3 Genome sequencing, assembly and annotation

The genomic DNA of 294 Roseobacter genomes was extracted using the Bacterial Genome DNA Rapid Extraction Kit (Guangzhou Dongsheng Biotech Co., Ltd.) and sequenced at the Beijing Genomics Institute (BGI, China) using the Illumina Hiseq xten PE150 platform. Raw reads were quality trimmed using Trimmomatic v0.39 (*62*) with options ‘SLIDINGWINDOW:4:15 MAXINFO:40:0.9 MINLEN:40’ and assembled using SPAdes v3.10.1(*63*) with ‘-careful’ option. Only contigs with length > 1,000 bp and sequencing depth > 5x were retained. Genome completeness, contamination, and strain heterogeneity (Table S3) were calculated using CheckM v1.1.2 (*64*).

Six isolates (SM26-46, SC33-45, SC33-90, MC52-69, SC1-11 and SC7-37) were additionally sequenced with the Nanopore platform (Nextomics Biosciences Co., Ltd.) to retrieve complete and closed genomes. The mismatches between Nanopore and Illumina reads were reconciled according to the following procedure. Raw reads of the Nanopore sequencing were first corrected by Necat v0.0.1 (*65*) with ‘PREP_OUTPUT_COVERAGE=100 CNS_OUTPUT_COVERAGE=50’ parameters. The polished reads were then assembled using Flye v2.6 (*66*) with the ‘--plasmids’ parameter. The initial assemblies were corrected twice using the polished Nanopore sequencing reads by Racon v1.4.13 (*67*) with ‘-m 8 -x -6 -g -8 -w 500’ options and five times using the Illumina sequencing reads by Pilon v1.23 (*68*) with default parameters. The Bandage v0.8.1 (*69*) was used to check whether the final assembled chromosomes and plasmids were closed, which showed that the chromosome and plasmids in all six genomes are closed except the plasmid 2b_P2 in MC52-69 and plasmid 2b_P5 in SC33-90 (Table S4).

Protein-coding genes were identified using Prokka v1.14.6 (*70*) with default parameters, and their functions were annotated using online RAST (*71*) and KEGG server (*72*). Genomic islands were predicted using Alien_hunter v1.7 (*73*) with default parameters.

#### 1.4 Phylogenomic tree construction

An ML phylogenetic tree was constructed based on 120 conserved bacterial genes (*74*) at the amino acid level to identify the phylogenetic position of 294 sequenced Roseobacter genomes. Other reference Roseobacter genomes included in the phylogeny were used following previous studies (*35, 75*). The 120 conserved proteins each were aligned using MAFFT v7.222 (*76*) with default parameters and trimmed using trimAl with ‘-resoverlap 0.55 -seqoverlap 60’ options. The trimmed alignments were linked together to form a super-alignment for each genome. The phylogenetic tree was constructed using IQTREE v1.6.12 (*57*) with the Modelfinder (*58*) for model selection, and a total of 1,000 ultrafast bootstrap replicates were sampled to assess the robustness of the phylogeny (*59*). The phylogenetic tree was visualized using iTOL v5.6 (*60*).

#### 1.5 Plasmid identification for the clade-2e1 members

Since a closed genome was not available to the clade-2e1, we used the following procedure to detect whether a contig is part of the chromosome or plasmid. Contigs of clade-2e1 genomes were aligned to the complete and closed genome of clade-2e2 (SC7-37) using Parsnp v1.2 (*77*). The contig was considered to be located on the chromosome or plasmid if > 80% region of this contig was aligned to the chromosome or plasmid of SC7-37. The remaining contigs were considered as unassigned.

#### 1.6 Population genomics analyses

The whole-genome average nucleotide identity (ANI) was calculated using FastANI v1.3 (*78*) to assess the genomic sequence similarity within and between clades. Besides, single nucleotide polymorphisms (SNPs) were identified using Parsnp v1.2 (*77*) with default parameters, and the SNP density was calculated using 10 Kbp sliding windows and plotted with custom scripts in R v3.6.1 (*79*). To investigate the genomic structural variation between clades, two complete and closed genomes (SC1-11 in clade-2d and SC7-37 in clade-2e2) were aligned using Mauve v2015-02-26 (*80*) with default parameters and the segment arrangement of the chromosome and the plasmid was visualized using Mauve with the ‘Min LCB weight’ parameter around 2,000. A similar comparison was also performed for two complete and closed genomes from clade-2a (SM26-46) and clade-2b (SC33-45), and for three complete and closed genomes within clade-2b (SC33-45, SC33-90 and MC52-69), respectively.

The clustering of accessory genes within clade was used to investigate whether the association with different hosts caused the differentiation of roseobacters at the genome content level. Orthologous gene families (OGs) were predicted using Roary v3.13.0 (*81*) with default parameters for genomes of the eight clades separately. The presence/absence pattern of accessory OGs was summarized as a binary matrix. The Euclidean distance of each pair of genomes was calculated using TBtools v1.0695 (*82*), and then genomes were clustered with the ‘complete’ method and visualized using TBtools.

#### 1.7 Co-cultivation of an axenic diatom culture and the Roseobacter isolates

The 11 Roseobacter representatives each were co-cultured with an axenic diatom culture *Phaeodactylum tricornutum* CCMP2561 to verify the effect of these roseobacters on the microalgal growth. The axenic microalgal culture was obtained from the Institute of Hydrobiology, Chinese Academy of Sciences, Wuhan, China. The diatom was inoculated in axenic F/2 medium at 20°C with 12 h/12 h light-dark cycle at 200 μmol photons m^-2^ s^-1^. The microalgal cells were counted three times each day using the Beckman Coulter Z2 (Beckman Coulter Inc., America) until they reached the logarithmic growth phase. The initial cell concentration of the microalgae used for growth assay was about 10^6^ cells mL^-1^.

The 11 Roseobacter strains each were inoculated in 5 mL of 2216E liquid medium and incubated at 25°C with 150 rpm shaking until the OD_600_ reached 0.6 - 0.7. For each strain, 3 mL of bacterial culture was centrifuged for 1 min at 12,000 rpm, and pellets were washed three times with sterile seawater and re-suspended in 100 μL of sterile seawater.

The 100 μL of bacterial suspension was inoculated in 30 mL of axenic microalgal culture as the experimental group, and another 30 mL of the axenic microalgal culture without bacterial inoculations were used as the negative control. Three replicates were set for all control and experimental groups. The microalgal cells were counted in triplicate at the third day of the co-culture experiment. The specific growth rate of the microalgae (μ_c_ for the control group and μ_e_ for the experimental groups) over three days was calculated as:

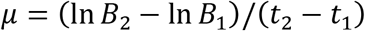

where B_1_ and B_2_ were the cellular density (concentration) in the culture at t_1_ (0 d) and t_2_ (3 d), respectively (*83, 84*). The significance level between μ_c_ and μ_e_ was estimated using the Student’s t-test.

#### 1.8 Phylogenetic analysis for flagellar gene clusters

Three homologous and phylogenetically distinguishable types of flagellar gene cluster (FGC) have been identified in the Roseobacter group (*35*). While none of them were identified in closely related clade-2a and clade-2b members, its homologs were found in clade-2d, clade-2e1 and clade-2e2 members. Next, phylogenetic analysis was performed for all the 83 genomes from the latter three clades to identify their FGC type. Four flagellar marker genes (*fliF, flgH, flgI, and flhA*) (*35*) were aligned at the amino acid level using MAFFT v7.222 (*76*) with default parameters and trimmed using trimAl with ‘-resoverlap 0.55 -seqoverlap 60’ options. The trimmed alignments were concatenated and a phylogenetic tree was constructed using IQTREE v1.6.12 (*57*) with the Modelfinder (*58*) function. A total of 1,000 ultrafast bootstrap replicates were sampled to assess the robustness of the phylogeny (*59*), and the phylogenetic tree was visualized using iTOL v5.6 (*60*).

#### 1.9 Motility assay

Eleven roseobacters were used in the motility assay, including three strains in clade-2a, three in clade-2b, two in clade-2d, and three in clade-2e2. The clade-2e1 was not included because all strains in this clade were lost. The 11 strains each were inoculated in 5 mL of 2216E liquid medium and incubated at 25°C with 150 rpm shaking until the optical density value at 600 nm (OD_600_) reached 0.6-0.7. The flagella and pili were observed under a transmission electron microscope (TEM; JEM-F200, Japan), and the motility of representative strains was tested on solid and liquid medium.

Aliquots (6 mL) of liquid culture for each bacterial strain was centrifuged at 1,000 rpm for 10 min. The pellets were rinsed with axenic water 2-3 times. Pellets were fixed overnight with 1 mL of 2.5% glutaraldehyde solution at 4°C. Fixed cells were resuspended with axenic water and stabilized on the 200-mesh copper grid (Beijing Zhongjingkeyi Technology Co., Ltd) with the carbon support film. Stabilized cells were stained with 2% sodium phosphotungstate solution for 1-2 min and observed by TEM.

The motility of the 11 strains were tested on soft agar plates with 0.2% and 0.3% agar (w/v) following previously study (*34, 85*). Soft agar plates were point inoculated with 3 μL of bacterial pre-cultures and incubated at 25°C for 6 d. The motility was also tested using sedimentation assays by inoculating 100 μL of bacterial cultures in 5 mL of fresh 2216E liquid medium and incubated at room temperature for 24 h without shaking (*86*).

### Text 2. Supplemental result

#### High genetic similarities of within-clade members are not due to clonal replication during laboratory cultivation

Considering that all members of each clade showed extremely high genetic identity at the DNA sequence level, it is possible that the collected isolates can be a mix of cells each representing a distinct genotype originally inhabiting the wild environment and cells replicated clonally during the laboratory cultivation of the microalgae. The latter situation needs to be considered since roseobacter isolation was performed after the one-month incubation of microalgae and since roseobacters divide approximately once per day (*87*). It is important to differentiate these two scenarios because the inference of population structure could be biased if laboratory clones dominated our collections. Here, we provide several lines of evidence against the laboratory replication hypothesis.

If all members of a clade would have been laboratory replicates from a single ancestor, we asked whether the observed SNPs could be explained by mutations during the period of laboratory cultivation of the diatoms. If a simple assumption is made that each SNP site was caused by a mutation, then the expected number of mutations (*S*) can be estimated based on the spontaneous mutation rate (*μ*), bacterial growth rate or number of cell divisions per day (*G*), the alignment length (*L*), laboratory cultivation time (*T*) and number of genomes under comparison (*N*), according to the following equation:

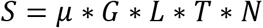

To make it simple, we used the mutation rate of the model Roseobacter strain, *Ruegeria pomeroyi* DSS-3 (*μ* = 1.39 × 10^−10^ per nucleotide per generation) (*88*) and the doubling time of wild roseobacters (*G* = *approximately one generation per day*) (*87*). Given that *L* = 4.34, 4.84, 4.37, 3.89, 3.56, 3.87, 3.74 and 4.31 Mb, and *N* = 29, 15, 100, 32, 16, 19, 48 and 35 genomes, for clade-1, clade-2a, clade-2b, clade-2c, clade-2d, clade-2e1, clade-2e2 and clade-3, respectively, the expected number of mutations of these clades are 0.53, 0.30, 1.82, 0.52, 0.24, 0.31, 0.75 and 0.63, respectively, under 30 days of laboratory cultivation of the diatoms. Since the expected numbers of mutations are all below one except clade-2b but there are a few dozen SNPs in six of the eight clades (Table S5), the observed SNPs cannot arise from mutations during the laboratory cultivation period.

One may argue that some of the SNPs could result from sequencing error. While it is difficult to distinguish between mutations and sequencing errors for singleton SNPs, presence of non-singleton SNPs (the rare variant found in at least two genomes) is convincing evidence against sequencing errors owing to the extremely low chance of having sequencing errors at the same sites. We identified non-singleton SNPs in six out of the eight clades (Table S5). In particular, all sequenced genomes from clade-1 and clade-2a each exhibited a unique combination of the non-singleton SNPs, suggesting that none of them represent laboratory clones (Table S5). For members from clade-2b, clade-2e1, clade-2e2 and clade-3, we identified 26, 14, 9 and 14 unique combinations of the non-singleton SNPs in 100, 19, 48 and 35 genomes, respectively, suggesting that at least a sizable number of the members in these clades represent the wild genotypes. In the case of the remaining two clades, clade-2c and clade-2d, no non-singleton SNPs were found in the 32 and 16 members, respectively.

It is important to note that the SNP analysis is restricted to the core DNA shared by all members of a clade, and members without displaying biologically meaningful SNPs in their core genomes may differ in their accessory genomes. We therefore asked whether genomes from each clade each harbor a unique combination of accessory genes (Table S5). However, this analysis is compromised by the incomplete genome assembly of reads derived from Illumina sequencing. This issue is mitigated, but not eradicated, when the genes missing in the complete and closed genome are used. For the four clades (clade-2a, clade-2b, clade-2d and clade-2e2) with at least one complete and closed genome, all of genomes each exhibit a unique combination of accessory genes (Table S5), which is evidence against the laboratory clonal replication hypothesis.

A higher-level marker to discriminate between genomes is gene order and genome rearrangement, which is best characterized using complete and closed genomes. We therefore closed three genomes (SC33-45, SC33-90 and MC52-69) from the same clade (clade-2b) each with additional sequencing by Nanopore (Table S4). Particularly, two of them (SC33-45 and SC33-90) are barely differentiated at the accessory genome content level (Fig. 2A), further highlighting the potential value of the genome rearrangement analysis. These three genomes (SC33-45, SC33-90 and MC52-69) differ in chromosomal DNA length (3,836,086 bp, 3,834,406 bp and 3,850,659 bp, respectively), and one DNA inversion event has occurred on their chromosome involving an orthologous DNA segment varying in their length (4644 bp, 4648 bp and 1164 bp, respectively; Fig. S2). They possess seven homologous plasmids, of which five are closed for all three genomes and thus are useful for comparison. While these five closed plasmids each have conserved gene order, they differ slightly in their lengths. Here are assembled lengths of these five closed plasmids in the three isolates: 2b_P1 (82,238 bp, 82,229 bp and 82,397 bp), 2b_P3 (290,592 bp, 290,600 bp and 290,590 bp), 2b_P4 (281,159 bp, 281,160 bp and 281,166 bp), 2b_P6 (210,913 bp, 210,913 bp and 210,915 bp), and 2b_P7 (110,566 bp, 110,566 bp and 110,565 bp).

Taken together, the available patterns of non-singleton SNPs in the core genomes, the strain-specific gene combination in the accessory genomes, the length of chromosomal DNA and plasmid DNA, and strain-specific order of genomic fragments strongly favor the hypothesis that most of the analyzed roseobacters each represents a distinct genotype originally from the wild environment.

**Fig. S1.**
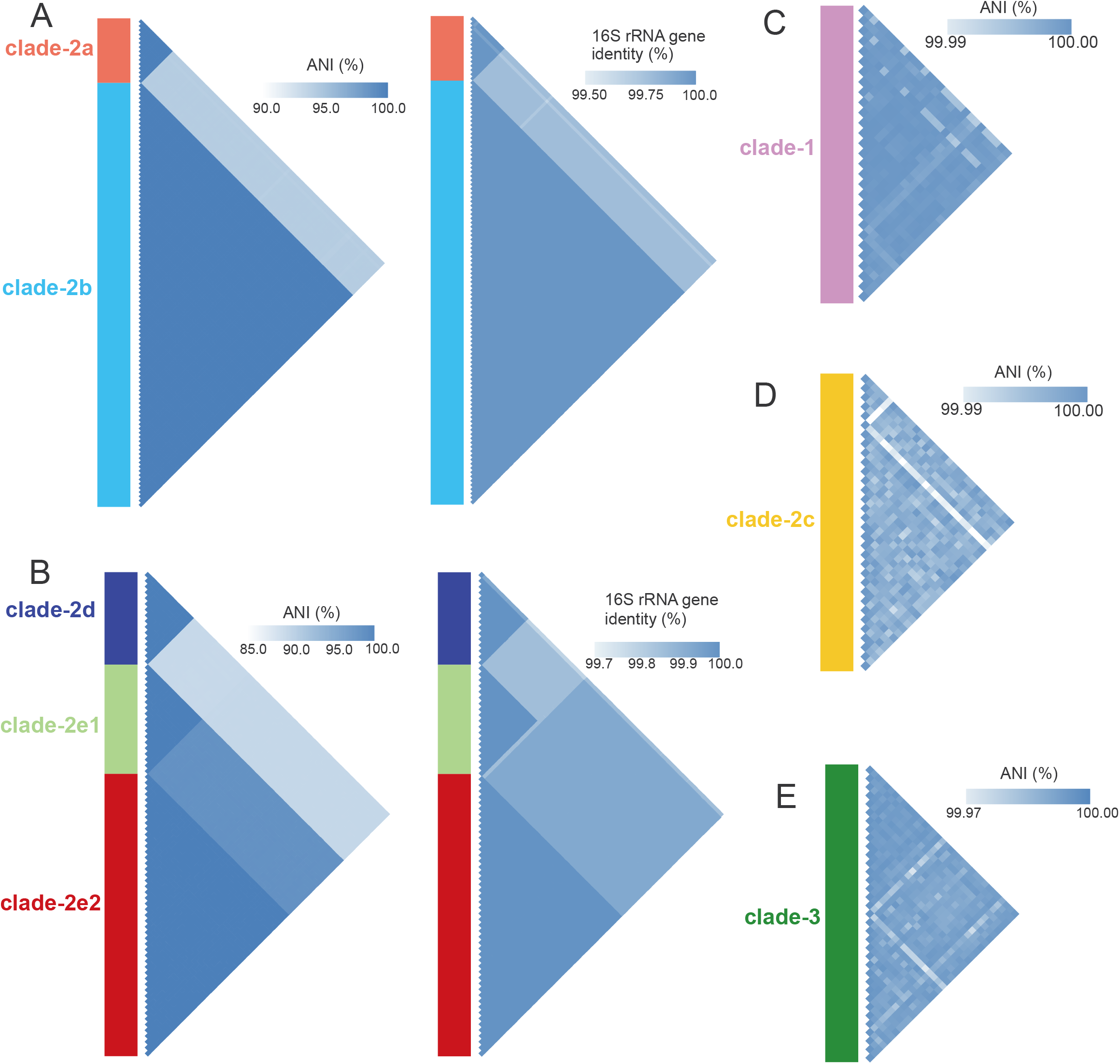
The whole-genome average nucleotide identity (ANI) and/or 16S rRNA gene identity for (A) clade-2a and clade-2b, (B) clade-2d, clade-2e1, and clade-2e2, (C) clade-1, (D) clade-2c, and (E) clade-3. The 16S rRNA gene identity is not shown for clade-1, clade-2c, and clade-3 because the 16S rRNA genes are identical within each clade.

**Fig. S2.**
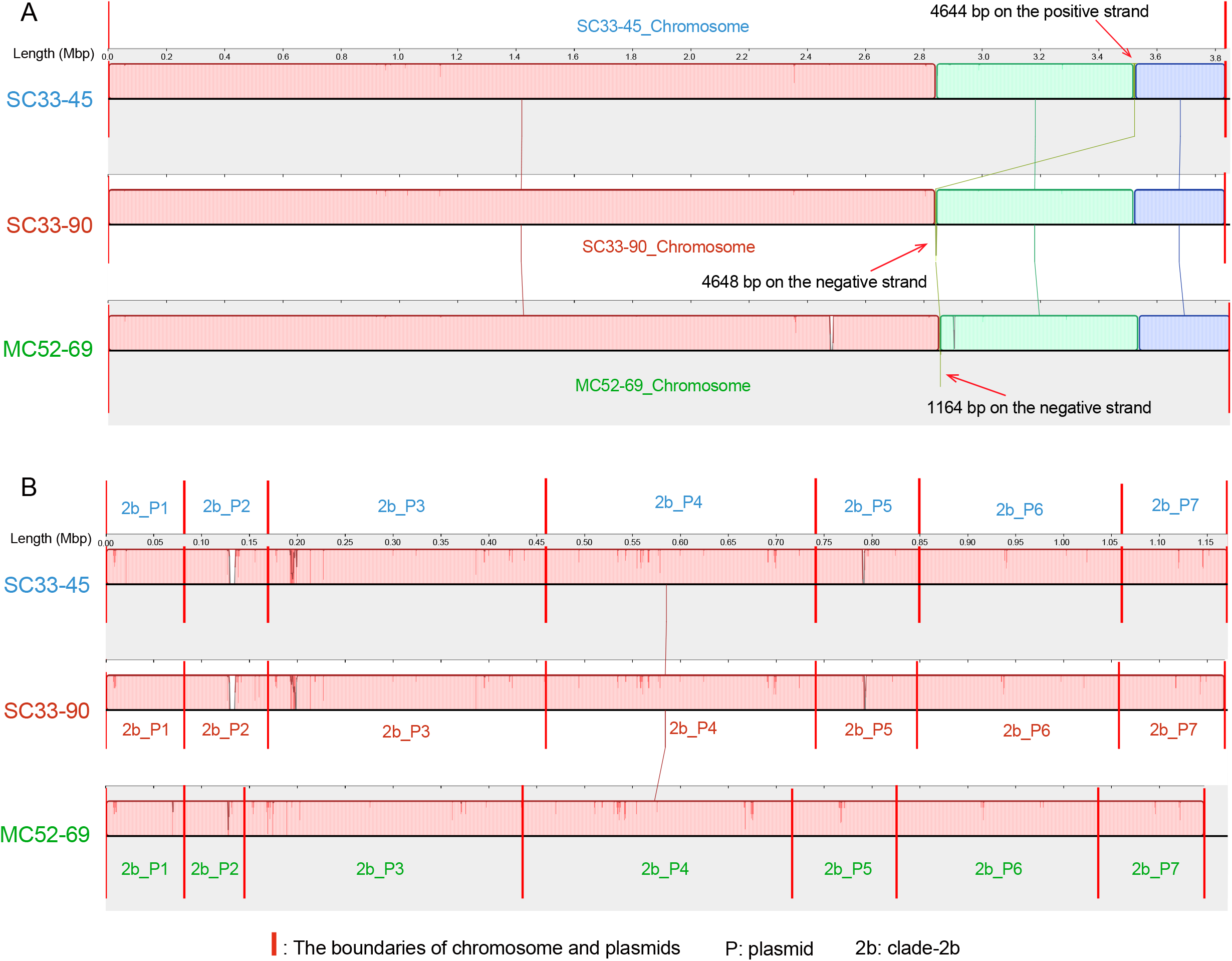
The genome arrangement of (A) chromosome and (B) plasmids of the three complete and closed genomes (MC52-69, SC33-45 and SC33-90) in clade-2b. Homologous regions shared by the three genomes are represented using locally collinear blocks (LCBs) with connected lines. The minimum LCB weight is 3485, which represents the minimum number of matching nucleotides identified in all LCBs. A similarity profile is shown within each LCB, and the height of the similarity profile represents the conservation level of the alignment. LCBs above and below the centerline represent genomic regions on the forward and reverse strand, respectively. The boundaries of replicons (chromosome and plasmids) are represented by red vertical lines.

**Fig. S3.**
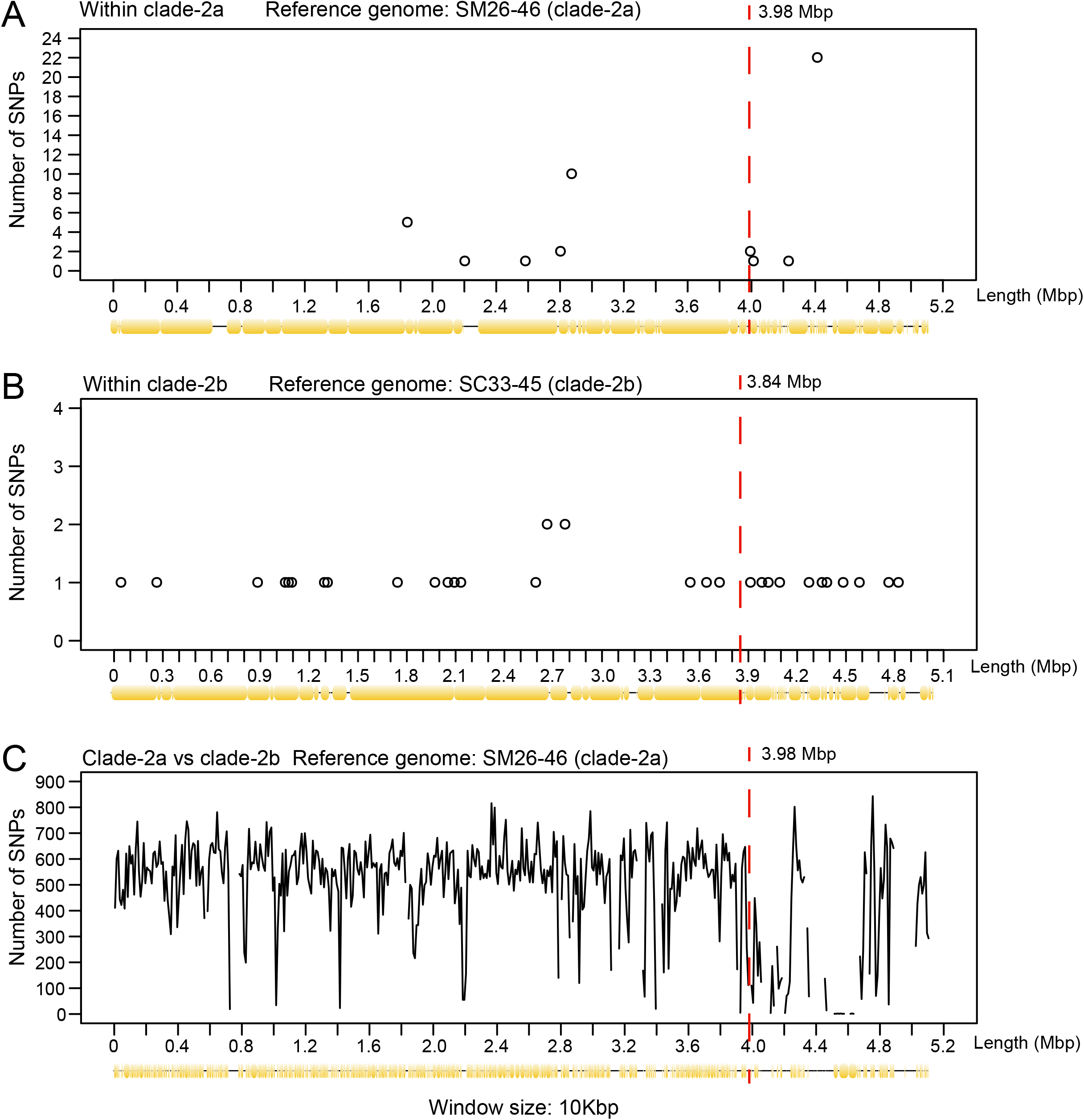
The density of non-singleton SNPs (A) within clade-2a, (B) within clade-2b, and (C) between these two clades. The SNP density was calculated with a window size of 10 Kbp. The aligned region is shown at the bottom of each panel. The boundary between chromosome and plasmid is shown using a red vertical dotted line.

**Fig. S4.**
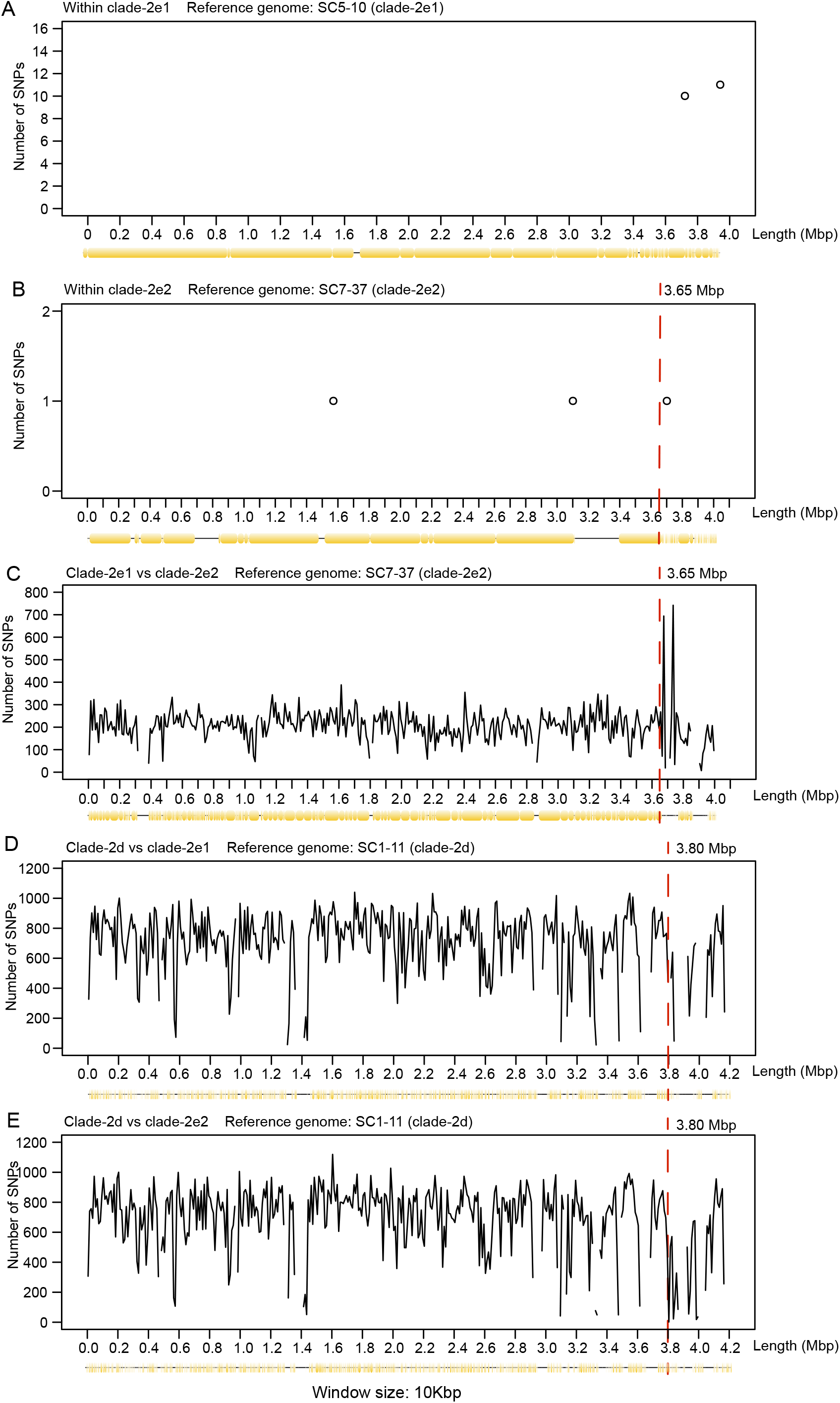
The density of non-singleton SNPs (A) within clade-2e1, (B) within clade-2e2, (C) between clade-2e1 and clade-2e2, (D) between clade-2d and clade-2e1, and (E) between clade-2d and clade-2e2. The SNP density of clade-2d is not shown because no SNP was found within this clade. The SNPs density was calculated with a window size of 10 Kbp. The aligned region was shown at the bottom of each panel. The boundary between chromosome and plasmid is shown using a red vertical dotted line.

**Fig. S5.**
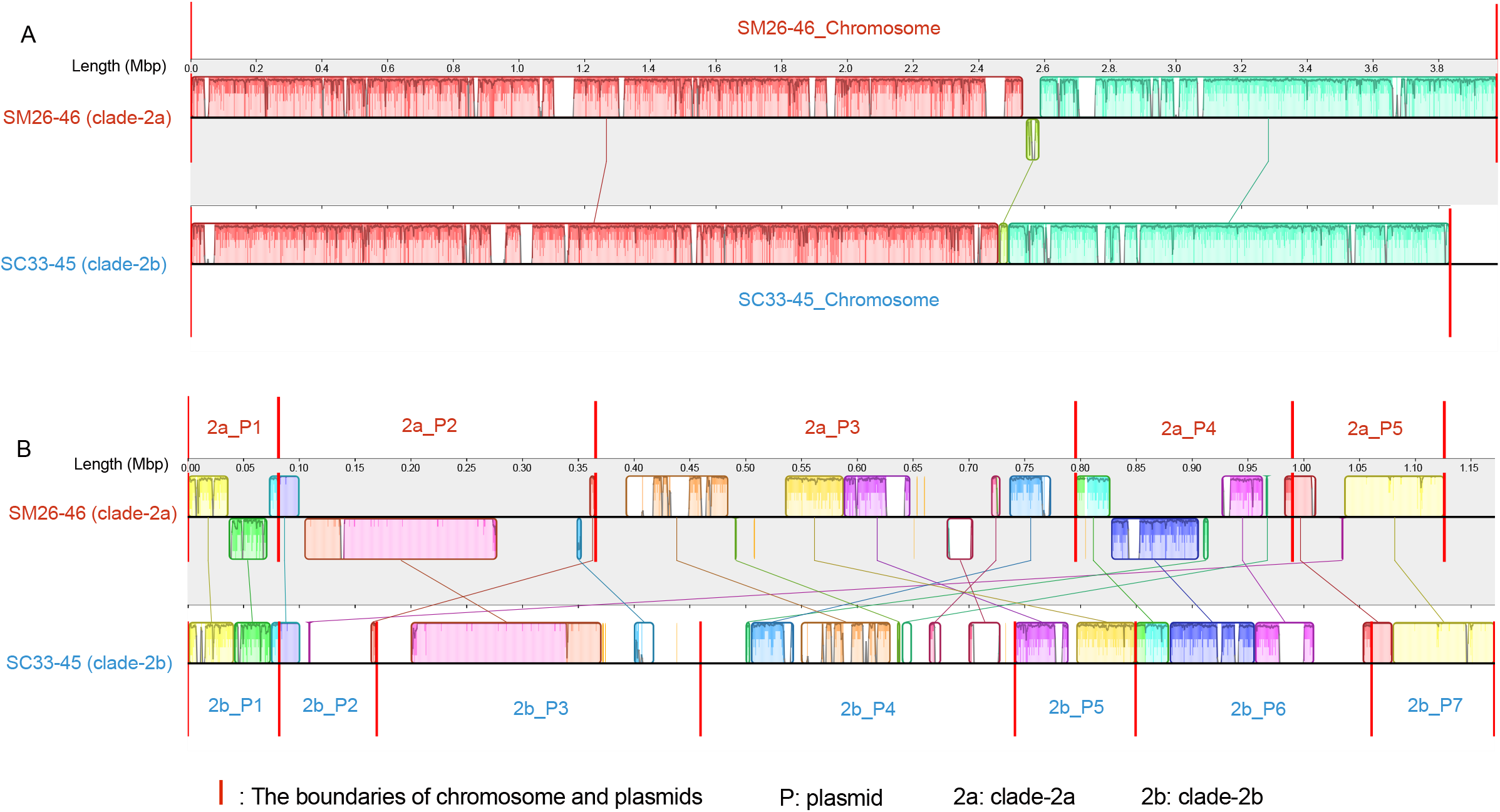
The genome arrangement of (A) chromosome and (B) plasmids of the two complete genomes in clade-2a (SM26-46) and clade-2b (SC33-45). Homologous regions shared by the two genomes are represented using locally collinear blocks (LCBs) with connected lines. The minimum LCB weight is 2249, which represents the minimum number of matching nucleotides identified in all LCBs. A similarity profile is shown within each LCB, and the height of the similarity profile represents the conservation level of the alignment. LCBs above and below the centerline represent genomic regions on the forward and reverse strand, respectively. The boundaries of replicons (chromosome and plasmids) are represented by red vertical lines.

**Fig. S6.**
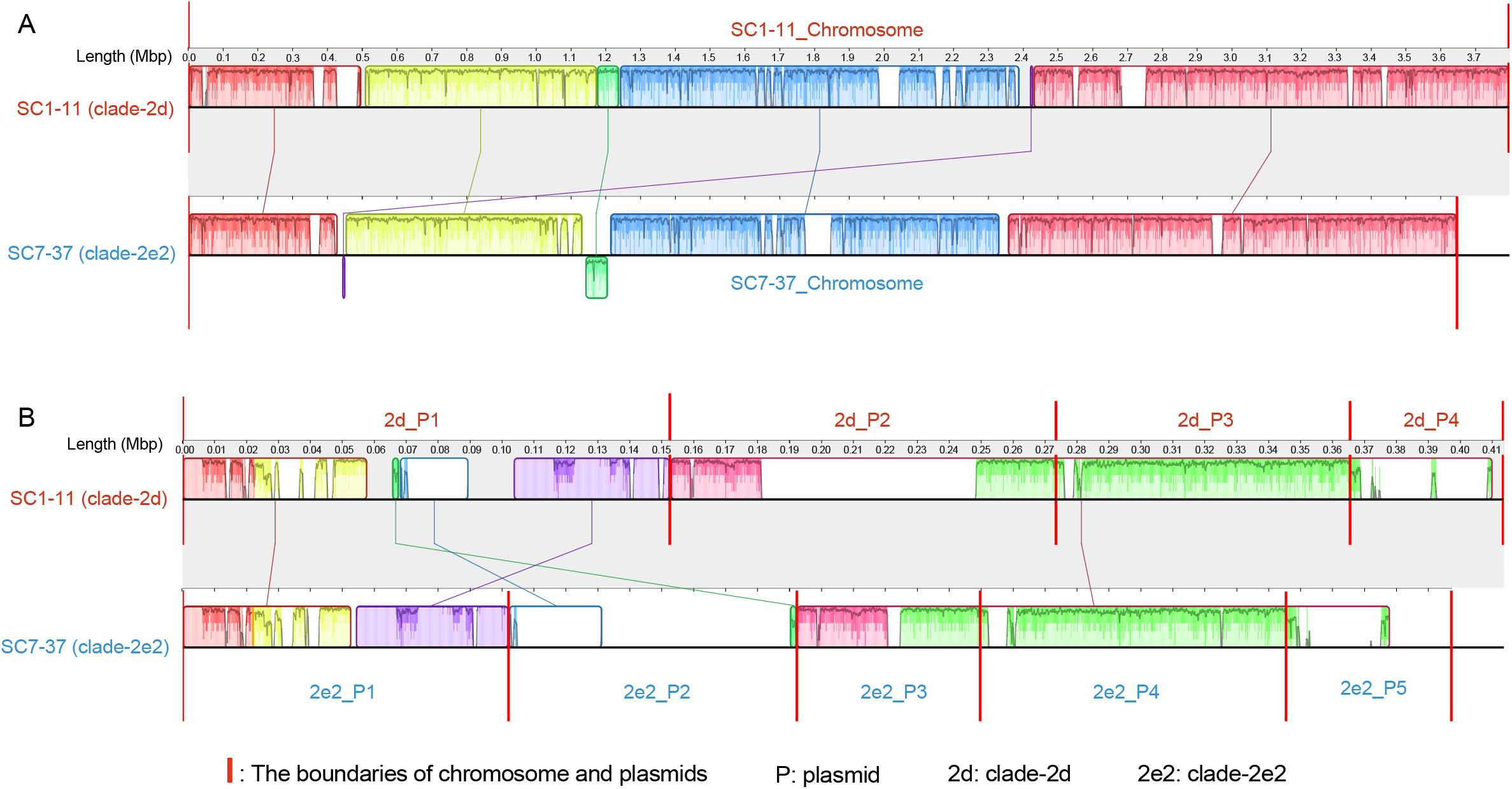
The genome arrangement of (A) chromosome and (B) plasmids of the two complete genomes from clade-2d (SC1-11) and clade-2e2 (SC7-37). Homologous regions shared by the two genomes are represented using locally collinear blocks (LCBs) with connected lines. The minimum LCB weight is 2209, which represents the minimum number of matching nucleotides identified in all LCBs. A similarity profile is shown within each LCB, and the height of the similarity profile represents the conservation level of the alignment. LCBs above and below the centerline represent genomic regions on the forward and reverse strand, respectively. The boundaries of replicons (chromosome and plasmids) are represented by red vertical lines.

**Fig. S7.**
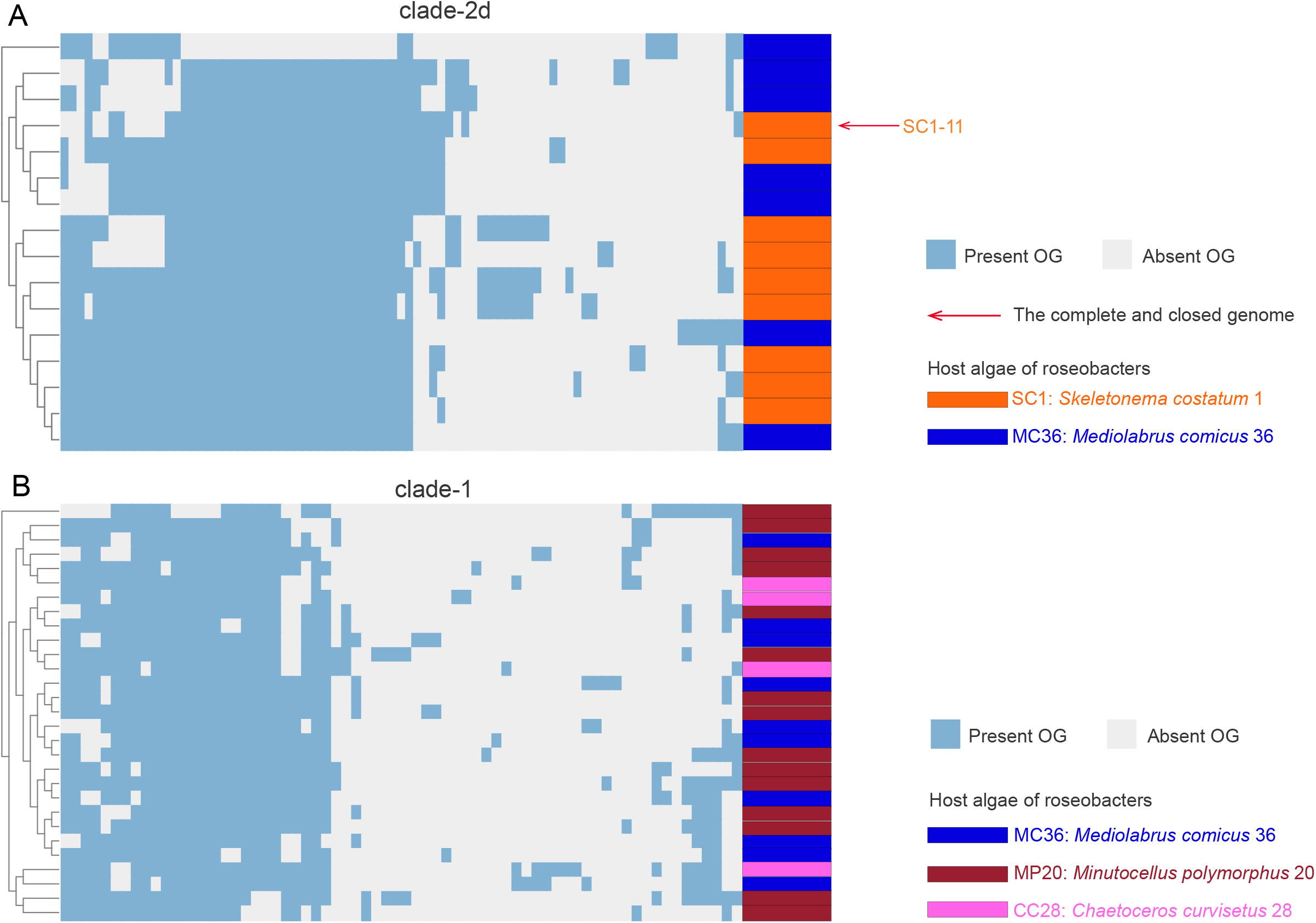
The clustering of accessory genes in the genomes of clade-2d (A) and clade-1 (B). The dendrogram of genome clustering was generated based on the presence and absence of their orthologous gene families (OGs), which are colored in blue and gray, respectively. The associated microalgae of bacterial strains are differentiated with colors. The complete and closed genome is marked with a red arrow.

**Fig. S8.**
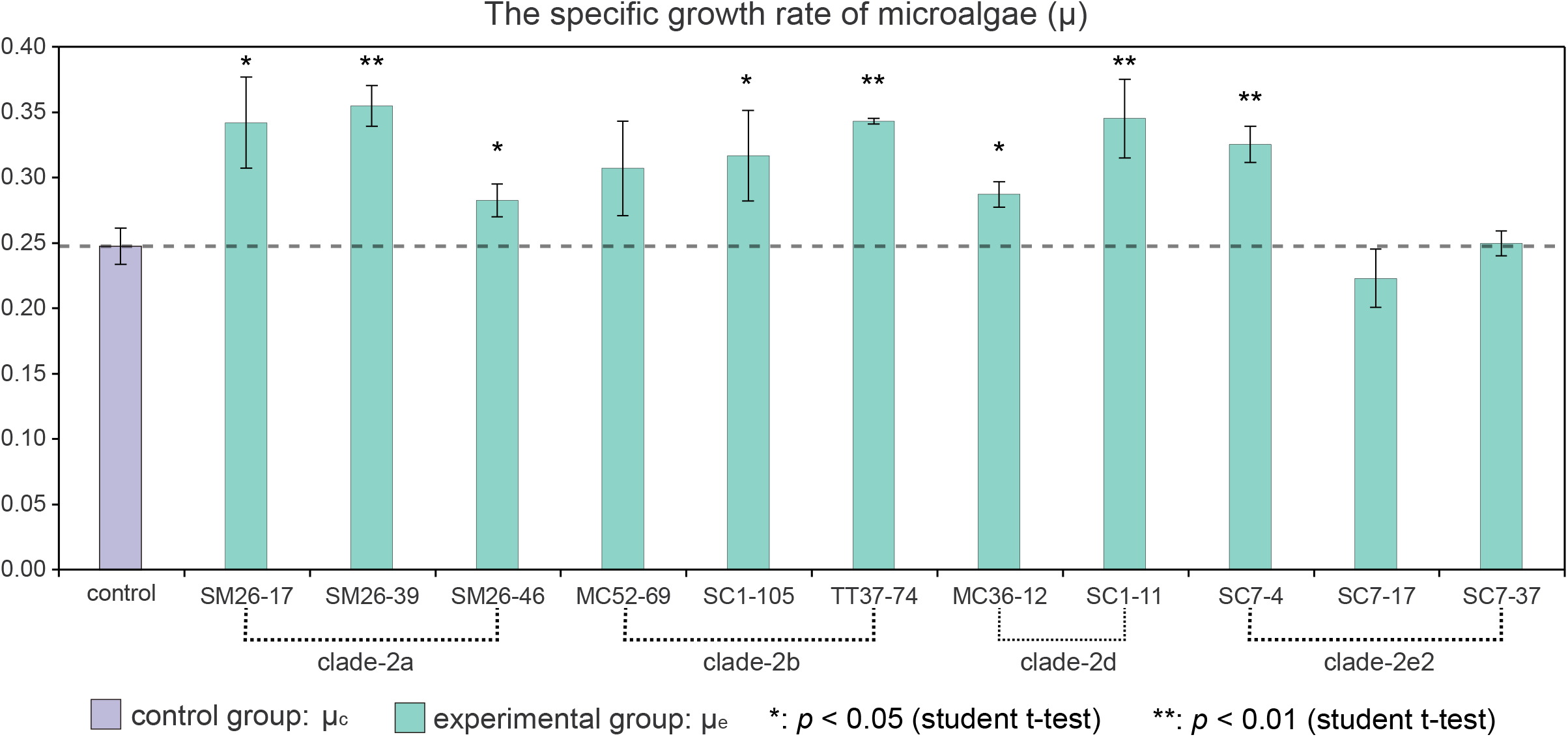
The specific growth rates of microalgal strains after three days of co-culture. The microalgal specific growth rate in the control groups (μ_c_) and experimental groups (μ_e_) are shown in purple and green columns, respectively. The significance level *p* < 0.05 and *p* < 0.01 compared to the control group is shown using * and **, respectively.

